# Connexin 43 impacts the chick premigratory cranial neural crest cell population without affecting the neural crest cell epithelial-to-mesenchymal transition

**DOI:** 10.1101/673921

**Authors:** Karyn Jourdeuil, Lisa A. Taneyhill

## Abstract

Gap junctions are intercellular channels that allow for the diffusion of small ions and solutes between coupled cells. Connexin 43 (Cx43), also known as Gap Junction Protein α1, is the most broadly expressed gap junction protein in vertebrate development. Cx43 is strongly expressed in premigratory cranial neural crest cells and is maintained throughout the neural crest cell epithelial-to-mesenchymal transition (EMT), but its function in these cells is not known. To this end, we have used a combination of *in vivo* and *ex vivo* live imaging with confocal microscopy, immunohistochemistry, and functional assays to assess gap junction formation, and Cx43 function, in chick premigratory cranial neural crest cells. Our results demonstrate that gap junctions exist between chick premigratory and migratory cranial neural crest cells, with Cx43 depletion inhibiting the function of gap junctions. While a reduction in Cx43 levels just prior to neural crest cell EMT did not affect EMT and subsequent emigration of neural crest cells from the neural tube, the size of the premigratory neural crest cell domain was decreased in the absence of any changes in cell proliferation or death. Collectively, these data identify a role for Cx43 within the chick premigratory cranial neural crest cell population prior to EMT and migration.

## INTRODUCTION

Connexins are integral building blocks of gap junctions (Li *et al.*, 2002; Laird, 2014; Sorgen *et al.*, 2018), which are intercellular channels that permit diffusion of small (<1 kDa) molecules and ions between cells (Goodenough *et al.*, 1996; Mese *et al.*, 2007; Abbaci *et al.*, 2008; Defranco *et al.*, 2008; Goodenough & Paul, 2009; Delmar *et al.*, 2018). Connexin proteins have four membrane spanning domains, two extracellular loops, one intracellular loop, and intracellular amino and carboxyl termini (Mese *et al.*, 2007; Abbaci *et al.*, 2008; Defranco *et al.*, 2008; Delmar *et al.*, 2018). Following their synthesis, six connexin proteins are assembled into a hemichannel, also known as a connexon, which can be either homomeric or heteromeric (Goodenough *et al.*, 1996; Mese *et al.*, 2007; Abbaci *et al.*, 2008; Defranco *et al.*, 2008; Goodenough & Paul, 2009; Delmar *et al.*, 2018; Solan & Lampe, 2018). To form a gap junction, hemichannels are trafficked to the cell membrane where they dock head-to-head with a hemichannel/connexon on an adjacent cell, in a homomeric or heteromeric manner, with the newly formed gap junction possessing a half-life of only a few hours (Mese *et al.*, 2007; Dbouk *et al.*, 2009; Goodenough & Paul, 2009). Notably, the connexin composition of gap junctions endows the gap junction with distinct properties including pore side, voltage-dependent gating, open probability, and permeability (Vogel & Weingart, 2002; Kanaporis *et al.*, 2011). More recently, connexin proteins have been discovered to have gap junction-independent roles, including binding and organizing cytoskeletal and signaling components as well as regulating junctional conductance and voltage sensitivity (Dbouk *et al.*, 2009; Delmar *et al.*, 2018; Sorgen *et al.*, 2018). Not surprisingly, connexins have been implicated in many human diseases involving a variety of tissues, including cancer metastasis, although cancer preventative roles have also been ascribed to gap junctions in certain tissue types, highlighting the dynamic nature of gap junction interactions and function (Davy *et al.*, 2006; Laird, 2014; Delmar *et al.*, 2018; Cooreman *et al.*, 2019; Waning *et al.*, 2019; J. I. Wu & Wang, 2019).

In humans, 21 connexins have been identified and there are at least 16 in mice (Li *et al.*, 2002; Laird, 2014) and, of these, Connexin 43 (Cx43) in the most abundant and the best studied (Goodenough *et al.*, 1996; Laird, 2014; Delmar *et al.*, 2018; Sorgen *et al.*, 2018; J. I. Wu & Wang, 2019). Our recent work revealed that Cx43 is expressed in chick cranial neural crest cells, including both prior to and during their epithelial-to-mesenchymal transition (EMT), and their subsequent migration (Jourdeuil & Taneyhill, 2018). Neural crest cells are a multipotent cell population induced during early embryogenesis within the neural plate border, a region between the neural and non-neural ectoderm (Donoghue *et al.*, 2008; Sauka-Spengler & Bronner-Fraser, 2008; Gammill & Roffers-Agarwal, 2010; Bronner, 2012; Bronner & LeDouarin, 2012; Ivashkin & Adameyko, 2013; Simoes-Costa & Bronner, 2013). During neurulation, premigratory neural crest cells, which lie at the dorsal aspect of the neural folds, undergo EMT, delaminate, and migrate into the periphery where they give rise to a large number of derivatives including intramembranous bones, satellite glial cells, melanocytes, smooth muscle, adipocytes, and odontoblasts (Sauka-Spengler & Bronner-Fraser, 2008; Bronner, 2012; Bronner & LeDouarin, 2012; Ishii *et al.*, 2012; Prasad *et al.*, 2012; Stuhlmiller & Garcia-Castro, 2012; Ivashkin & Adameyko, 2013; Simoes-Costa & Bronner, 2013). While Cx43 (and gap junctions) has been identified in human, *Xenopus*, and mouse migratory neural crest cells (Reaume *et al.*, 1995; Ewart *et al.*, 1997; Lo *et al.*, 1997; Huang *et al.*, 1998; Waldo *et al.*, 1999; Xu *et al.*, 2001; Li *et al.*, 2002; Xu *et al.*, 2006; Rhee *et al.*, 2009; Francis *et al.*, 2011; Kotini *et al.*, 2018), the presence of functional gap junctions in the premigratory cranial neural crest cell population, however, remains unknown. Moreover, nearly 80 distinct mutations in *Cx43* gives rise to a multisystem developmental disorder called oculodentodigital dysplasia (ODDD), which is characterized by defects in neural crest-derived craniofacial bones (William A. Paznekas *et al.*, 2003; W. A. Paznekas et al., 2009; Laird, 2014; Delmar *et al.*, 2018). Gap junction-related disorders of the peripheral nervous system have also been identified, which, in the craniofacial region, are derived from neural crest and placode cells (Hamburger, 1961; Delmar *et al.*, 2018).

The robust expression of Cx43 in premigratory cranial neural crest cells, both before and during EMT, suggests that gap junctions may exist within this cell population to facilitate intercellular communication as these cells dismantle their adherens and tight junctions during EMT (Sauka-Spengler & Bronner-Fraser, 2008; Bronner, 2012; Schiffmacher *et al.*, 2014; Schiffmacher *et al.*, 2016; Schiffmacher *et al.*, 2018). A recent study in *Xenopus* cranial neural crest cells identified a 20 kDa isoform of Cx43 (Cx43-20k) that trans-activates *N-cadherin* to promote NCC migration without affecting the premigratory neural crest cell population or EMT (Kotini *et al.*, 2018). In chick, however, *N-cadherin* is extinguished very early within the premigratory cranial neural crest cell population and is not expressed by migratory cranial neural crest cells until later in development (Dady *et al.*, 2012). To further explore a role for gap junctions and Cx43 within the cranial neural crest cell population of the chick, we performed live imaging and loss-of-function assays. Our data reveal that functional gap junctions are formed between premigratory and migratory neural crest cells. In addition, depletion of Cx43 is sufficient to inhibit gap junction function in both premigratory and migratory neural crest cells, but this does not overtly affect neural crest cell EMT and early migration. Instead, a reduction in Cx43 resulted in a concomitant loss of Snail2- and Pax7-positive premigratory cranial neural crest cells. Altogether, these data suggest that Cx43 plays a role in influencing the size of the premigratory cranial neural crest cell population in the chick embryo, but, remarkably, Cx43-dependent gap junctions are likely not required for neural crest cell EMT, implying that role of Cx43, and gap junctions, may be unique to individual species.

## MATERIALS AND METHODS

### Chicken embryo culturing and fixation

Fertilized chicken eggs were obtained from both Moyer’s Chicks (Quakertown, PA, USA) as well as the University of Maryland (College Park, MD, USA) and incubated at 38°C in humidified incubators (EggCartons.com, Manchaug, MA, USA). After 30 hours of incubation, eggs were removed from the incubator and a window was made in the shell to allow access to the embryo. Samples were staged according to the Hamburger and Hamilton (Hamburger & Hamilton, 1951) staging table. Manipulations were performed between Hamburger and Hamilton (HH) stage 7^+^ through to HH8^−^ (2-3 somite stage). Embryos were subsequently collected between HH8^−^ and HH10 (3-10 somite stage).

### Cx43 and mismatch control morpholinos

A 3’ lissamine-tagged antisense translation-blocking Cx43 morpholino (Cx43MO, 5’- CAGGGCACTCCAATCACC<u>CAT</u>CTTC-3’), or a 5-base pair mismatch Cx43 control morpholino (Cx43MM, 5’-CAcGGCAaTCCAATaACa<u>CAT</u>aTTC-3’) (start codon underlined and mismatches shown in lower case), was designed to target the *Cx43* transcript according to the manufacturer’s criteria or serve as a control, respectively (Gene Tools, LLC, Philomath, OR). Both morpholinos were used at a concentration of 500 μM as previously described (Fishwick *et al.*, 2012; Schiffmacher *et al.*, 2014). We confirmed that the morpholinos were specific to *Cx43* and did not bind to any other sites by using the NCBI Nucleotide BLAST tool to search the invert complement of the morpholino sequence against the chicken genome, as per Gene Tools’ recommendations). Immunoblotting was also performed to demonstrate evidence of Cx43 knockdown (see below).

### Dorsoventral electroporation of the neural crest

Dorsoventral electroporation of the neural tube was used to target the entire midbrain premigratory cranial neural crest cell population between HH7^+^ and HH8^−^. To this end, a drop of Ringers Solution (pH 7.4) containing 0.1% Penicillin-Streptomycin (10,000 U/mL; Gibco, 15140122) was applied to the vitelline membrane overlying the embryo. A hole was then made at the edge of the area opaca using a 20-gauge needle. The Cx43 MO or Cx43 control MO was injected between the vitelline membrane and the embryo using fine glass needles to completely flood the area that will give rise to the head. The electrodes were then placed such that the anode was inserted into the hole at the border of the area opaca, underlying the morpholino-flooded epithelium, with the cathode barely touching the vitelline membrane. Three, 9 V, 50 ms electric pulses were applied with a 200 ms refractory period in between. Following electroporation, another drop of Ringers with 0.1% Penicillin-Streptomycin was applied to the surface of the vitelline membrane, the eggs were sealed with tape and parafilm, and re-incubated for three hours. Embryos were then screened for the presence of Cx43 MO or Cx43 control MO using a Zeiss Discovery v8 stereomicroscope, with successfully electroporated embryos fixed using 4% Paraformaldehyde (PFA; Fisher Scientific, 30525-89-4) at 4°C overnight.

### In ovo unilateral electroporations

Unilateral electroporation of the early chick neural tube was conducted to target premigratory neural crest cells in the midbrain. The Cx43 MO or Cx43 control MO was injected into the lumen of the neural tube of HH7^+^ and HH8^−^ (2 or 3 somites) chick embryos using fine glass needles. Electrodes were then placed on either side of the midbrain and two, 25 V, 30 ms electric pulses were applied to the midbrain with a 200 ms refractory period in between (Gemini System, BTX). Following electroporation, the eggs were sealed with tape and parafilm, and returned to the incubator for three to six hours. Embryos were then screened as described above and subsequently fixed using 4% PFA or were processed for subsequent experimental analysis as outlined below.

### Immunoblotting

To validate both Cx43 antibodies used in these experiments (Sigma Aldrich, C6219, raised to the carboxy terminus (CT); Abcam, ab78055, raised to the amino terminus (NT)), chick heads were collected from HH8 and 9 (5 somites through to 8 somites) wild-type embryos. Similarly, the knockdown efficiency of the Cx43 MO was evaluated by collecting and independently pooling the electroporated and contralateral control half neural tubes from the midbrains of Cx43 MO and Cx43 control MO embryos at the 7 somite stage. Samples were rinsed in Ringer’s Solution, centrifuged at 500 *g* for 5 minutes at 4°C, and then snap-frozen in liquid nitrogen. Protein extraction and immunoblotting were then performed as previously described (Schiffmacher *et al.*, 2014; Shah *et al.*, 2017; Schiffmacher *et al.*, 2018). Briefly, cell pellets were lysed in lysis buffer (50 mM Tris-HCl pH 8.0, 150 mM NaCl, 1% IGEPAL CA-630, 1mM EDTA) supplemented with cOmplete Protease Inhibitor Cocktail (Sigma, 1167498001) and 1 mM PMSF (Sigma Aldrich, 10837091001) for 30 minutes at 4°C with periodic mixing. The clarified, solubilized protein fractions were collected following centrifugation at maximum speed for 15 minutes at 4°C and protein concentrations were quantified by Bradford assay (Thermo Fisher Scientific). Equivalent amounts of protein per sample were boiled at 99°C for five minutes in 4X reducing Laemmli sample buffer and then centrifuged at maximum speed for five minutes at room temperature. The samples were then loaded into a Novex Wedgewell Tris-Glycine gel (either 16% (wild-type; XP00160BOX, Invitrogen) or 10% (Cx43 MO and Cx43 control MO; XP00100BOX, Invitrogen)) and subsequently transferred to a 0.45 μm BioTrace PVDF membrane (Pall, Port Washington, NY). Membranes were incubated in blocking solution (5% dry milk in 1X PBS + 0.1% Tween-20 (PTW)) for one hour at room temperature and then incubated overnight at 4°C with the following primary antibodies diluted in blocking solution: CT Cx43 (1:1000 (wild-type); 1:5000 (Cx43 MO and Cx43 mismatch MO)) or NT Cx43 (1:1000). Membranes were washed in PTW and then incubated with species- and isotype-specific horseradish peroxidase-conjugated secondary antibodies (40 ng/ml; Jackson ImmunoResearch) in 5% blocking solution for one hour at room temperature. Membranes were washed again in PTW and antibody detection was performed using the Supersignal West Pico or Femto chemiluminescent substrates (Thermo Scientific) and visualized using a ChemiDoc XRS system (Bio-Rad). The immunoblots were then stripped using Restore Western Blot Stripping Buffer (Thermo Scientific) for two hours at 37°C and re-probed using a β-actin primary antibody (1:1000; Santa Cruz Biotechnology, sc-46668) and its species- and isotype-specific secondary antibody as described above. Immunoblots were analyzed using Image Lab software (Bio-Rad), in order to determine band size and volume. The volume of the Cx43 control MO- and Cx43 MO-electroporated (and their corresponding contralateral controls) were then normalized to the loading control in order to determine the efficacy of morpholino-mediated knockdown.

### Evaluation of gap junction function

Gap junction function was evaluated by introducing either the Cx43 MO or Cx43 control MO via dorsoventral electroporation of the premigratory cranial neural crest cell population as described. After three hours of incubation, the embryo, along with the extraembryonic membranes, was dissected off the yolk and carefully washed in Ringer’s Solution. The vitelline membrane was then removed and a ring of filter paper (VWR; 28310-128) was placed around the embryo so as to hold the embryo taut. A 1:1 solution of 1 mg/ml Calcein (C481; dissolved in Ringer’s Solution), which only passes to cells via gap junctions (Davy *et al.*, 2006; Abbaci *et al.*, 2008), and 1 mg/ml Dextran, Cascade Blue (D1976; dissolved in ddH_2_O), to allow for the identification of the injected cells as it does not pass through gap junctions (Ziambaras *et al.*, 1998), was injected into the dorsal neural folds using a fine glass needle. Excess Calcein-Dextran solution was washed away using Ringer’s Solution. Then, the embryo and filter paper were turned over and the tautness of the embryo and extraembryonic tissues were adjusted before a second ring of filter paper was applied to the back of the embryo, creating a sandwich. The embryo was then transferred, dorsal side down, onto a Glass Bottom Culture Dish (MatTek, P35G-1.5-20-C) with a 20 mm Microwell and covered with a mixture of 1% Low Melting Point Agarose (prepared with Ringer’s Solution; Promega, V2111) and thin albumin in order to immobilize the embryo on the dish.

To assess the effects of Cx43 knockdown on migratory neural crest cells, embryos were first unilaterally electroporated at HH7^+^ or HH8^−^ with either the Cx43 MO or Cx43 mismatch MO and then cultured for an additional three hours at 38°C. The electroporated embryos were dissected off the yolk and washed with Ringer’s Solution. The midbrain dorsal neural folds (containing the premigratory neural crest cells) were then carefully dissected from both the electroporated and the contralateral control side in PB-1 standard medium and placed into either a Glass Bottom Culture Dish with a 10 mm Microwell (MatTek, P35G-1.5-10-C (N=3)) or an 8-well Lab-Tek II Chamber Slide (Naige Nunc International, 154534; if the samples were to be immunostained (N=3)) that had been coated with Poly-L-Lysine (Sigma Aldrich, P5899) and Fibronectin (Corning, 356008) containing serum-free DMEM (Corning; 10-013-CV) supplemented with 0.1% Penicillin-Streptomycin and N-2 supplement (Gibco, 17501-048). To each of these treatments (Cx43 MO or Cx43 control MO), dissected wild-type neural folds that had been incubated for 30 minutes at 37°C in a 1:500 Calcein-AM (Invitrogen, C3099)/DMEM (supplemented with Penicillin-Streptomycin and N-2) solution in an uncoated chamber slide, and subsequently washed three times with DMEM, were added. The neural folds were then cultured overnight in order to allow the neural crest cells to undergo EMT and migrate out onto the underlying fibronectin. Cultures were then checked hourly until the Cx43 MO- or Cx43 control MO-treated neural crest cells made contact with the Calcein-AM-treated neural crest cells.

All cultures were imaged live using a Zeiss LSM800 Confocal with AiryScan detection held within a temperature-controlled unit which was maintained at 37°C and 5% CO_2_ for the duration of imaging. Images were acquired using Zen 2.0 (Blue Edition) and processed into videos using Adobe Photoshop CC 2018 and Adobe Premiere Pro CC 2018. After live imaging, samples that had been incubated in an 8-well chamber slide were fixed in 4% PFA for one hour at room temperature and prepared for immunostaining.

### Immunohistochemistry

After fixation, whole embryos were washed in 1X phosphate-buffered saline (PBS), gelatin-embedded, and serially sectioned at 16 μm. Samples were then de-gelatinized and blocked in 1X PBS containing 0.2% TritonX-100 (1X PBST; EMD Millipore Corporation, TX1561-1) and 10% sheep serum (Lampire, 733900). Cultured cells, on the other hand, were just washed in 1X PBST. Samples were then incubated with various primary antibodies (Cx43 (1:500; Sigma-Aldrich, C6219), Pax7 (1:10; Developmental Studies Hybridoma Bank (DSHB)), Sox10 (1:500; GeneTex, GTX128374), Snail2 (1:100; Cell Signaling Technology, C1967), Phospho-histone H3 (pHH3, 1:200; Millipore), N-cadherin (1:50; DSHB clone MNCD2), and E-cadherin (1:500; BD Transduction Laboratories, 610181)). The following day, after washing four times for 30 minutes with 1X PBST, sections were incubated with appropriate secondary antibodies and incubated for two to three hours at room temperature or overnight at 4°C (goat anti-rabbit IgG 488 or 647; Invitrogen, A11034 and Jackson ImmunoResearch, 111-605-003, respectively (Cx43 (1:500), Sox10 (1:500), pHH3 (1:200), and Snail2 (1:100)); goat anti-mouse IgG1 647; Invitrogen, A21240 (Pax7 (1:200) and E-cadherin (1:500)); goat anti-rat IgG; Life Technologies, A21247 (N-cadherin (1:200)). All primary and secondary antibodies were diluted in 1X PBST and 5% sheep serum at the indicated concentrations. After washing four times for 30 minutes, coverslips were mounted using DAPI Fluoromount-G (Southern Biotech, 0100-20) to mark cell nuclei. For all sectioned embryos, images of serial, transverse sections were acquired with the LSM Zeiss LSM800 confocal microscope with AiryScan detection and processed using Zen 2.0 (Blue Edition) software and Adobe Photoshop CC 2018.

### TUNEL

A TUNEL assay (*In situ* Cell Death Detection Kit, Fluorescein; Roche, 1164795910) was performed on 4% PFA-fixed, cryopreserved sections to detect apoptotic cells as described previously (C. Y. Wu *et al.*, 2014; Shah *et al.*, 2017). Following TUNEL, coverslips were mounted with DAPI Fluoromount-G and imaged as described above.

### Statistical analysis

The number of premigratory, migratory, and total neural crest cells were counted in at least five serial, transverse sections from Cx43 MO- or Cx43 control MO-electroporated dorsal neural folds and normalized to the number of premigratory, migratory, and total neural crest cells on the contralateral side, as in (Hutchins & Bronner, 2018) (see Table 1 for N values). Neural crest cells were identified with either Snail2, Pax7, or Sox10 immunostaining, as described above. A two-tailed Student’s t-test was then performed to assess statistical significance.

**Table 1:**
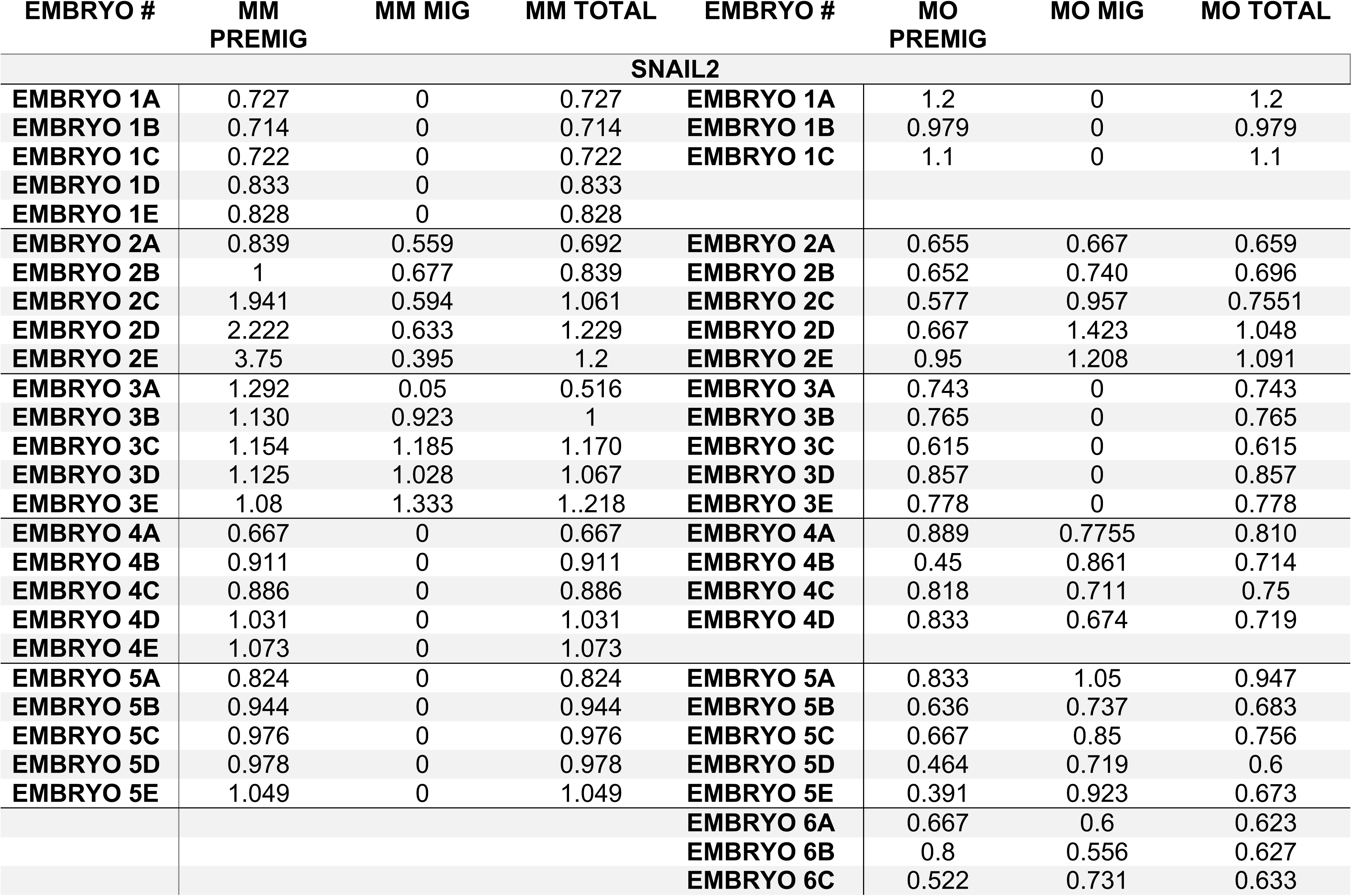
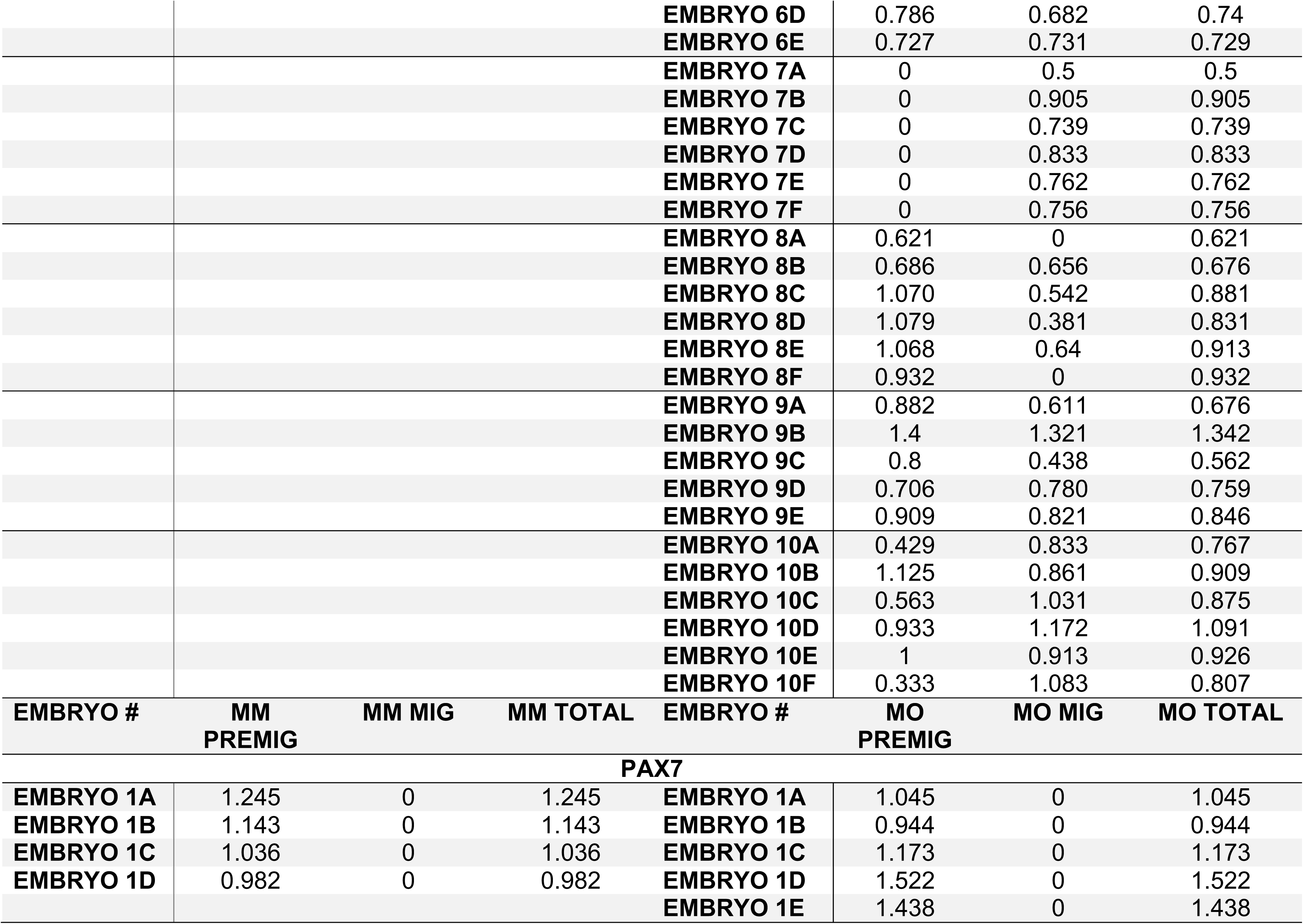
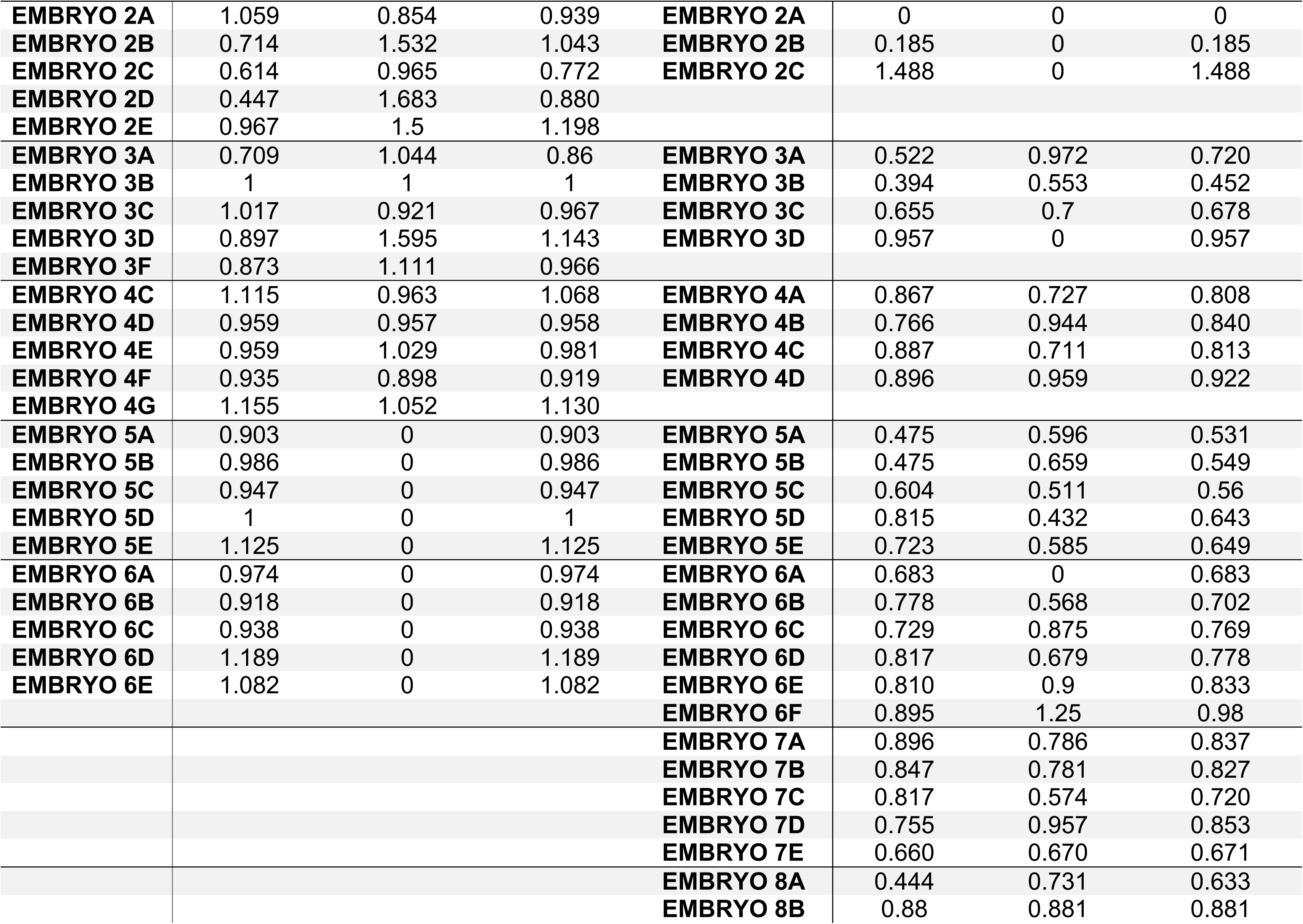
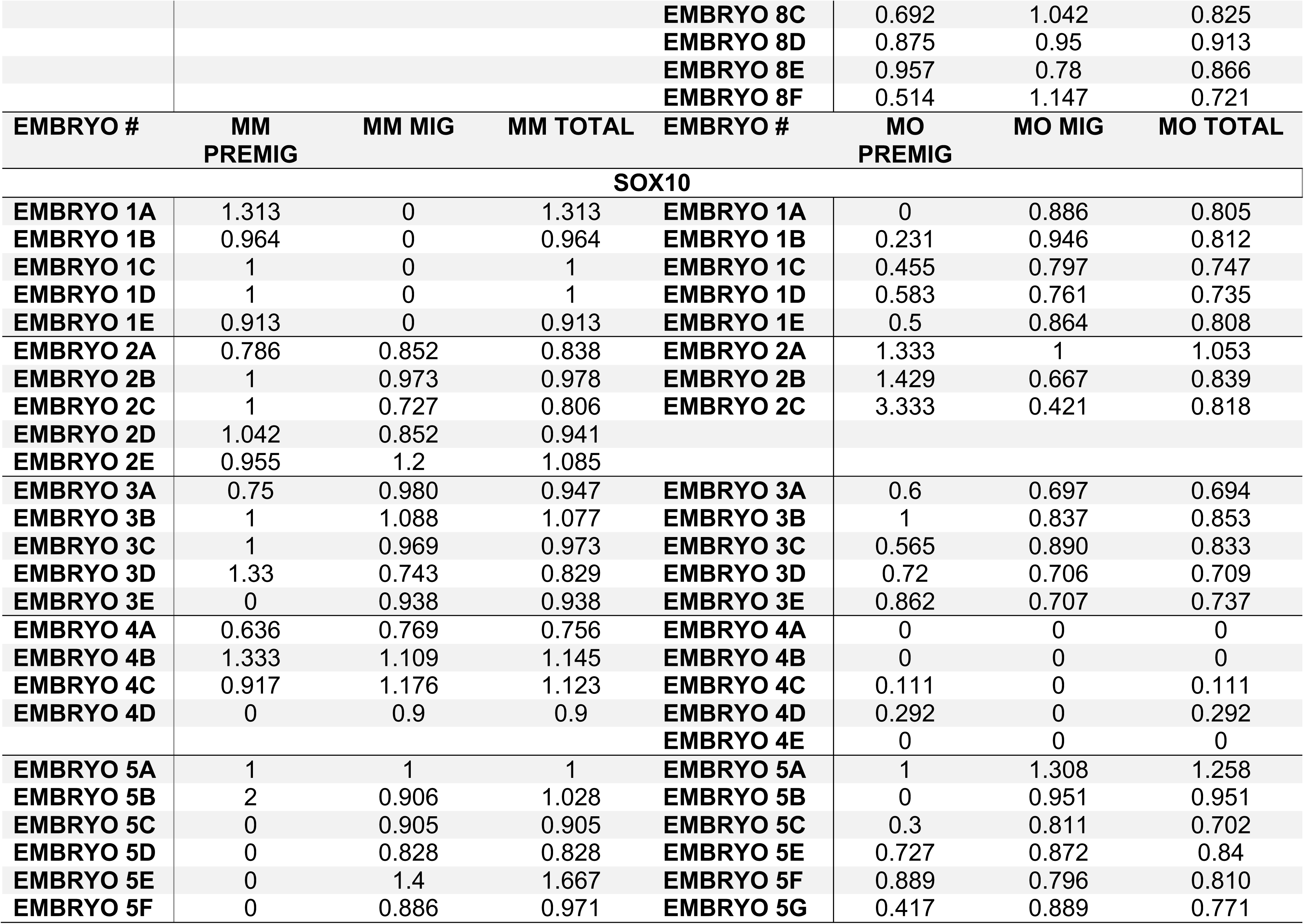
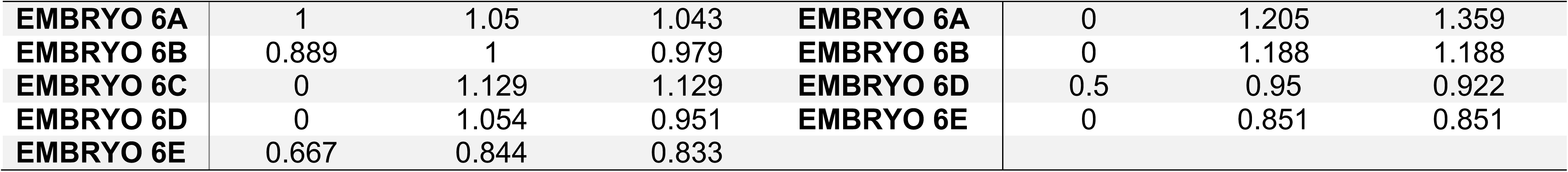
Cell counts for each embryo and section normalized to the contralateral, control side used for statistical analysis.

## RESULTS

### Multiple isoforms of Cx43 are present in the chick cranial neural crest

A recent study revealed a role for a short C-terminal isoform of Cx43 (Cx43-20k) during *Xenopus* cranial neural crest cell migration (Kotini *et al.*, 2018). To define the Cx43 isoforms expressed in the chick head prior to and during neural crest cell EMT and migration (HH8^+^ to HH9^+^ (5 somite stage (ss) to 8ss)), we performed immunoblotting for Cx43 using an antibody that recognizes the Cx43 C-terminus. These results revealed that only the full-length Cx43 isoform (37 kDa) is present in heads at both the 5ss, in the premigratory neural crest cell population, and at the 6ss, when neural crest cells first begin EMT (Fig. 1A). By the 7ss, when cranial neural crest cells are undergoing EMT, we noted a strong band corresponding to Cx43-20k and a lighter band of approximately 11 kDa, the latter of which was also identified previously but does not regulate *N-cadherin* expression (Kotini *et al.*, 2018) (Fig. 1A). The expression of the 20 and 11 kDa isoforms is also maintained at the 8ss (Fig. 1A). To confirm the identify of these bands, we performed immunoblotting for Cx43 using an antibody raised to the Cx43 N-terminus, which yielded a band of 37 kDa, corresponding to full-length Cx43 protein, and no C-terminal isoforms (Fig. 1B). These data point to a potential role of full-length Cx43 in premigratory neural crest cells and during early stages of neural crest cell EMT.

**Figure 1:**
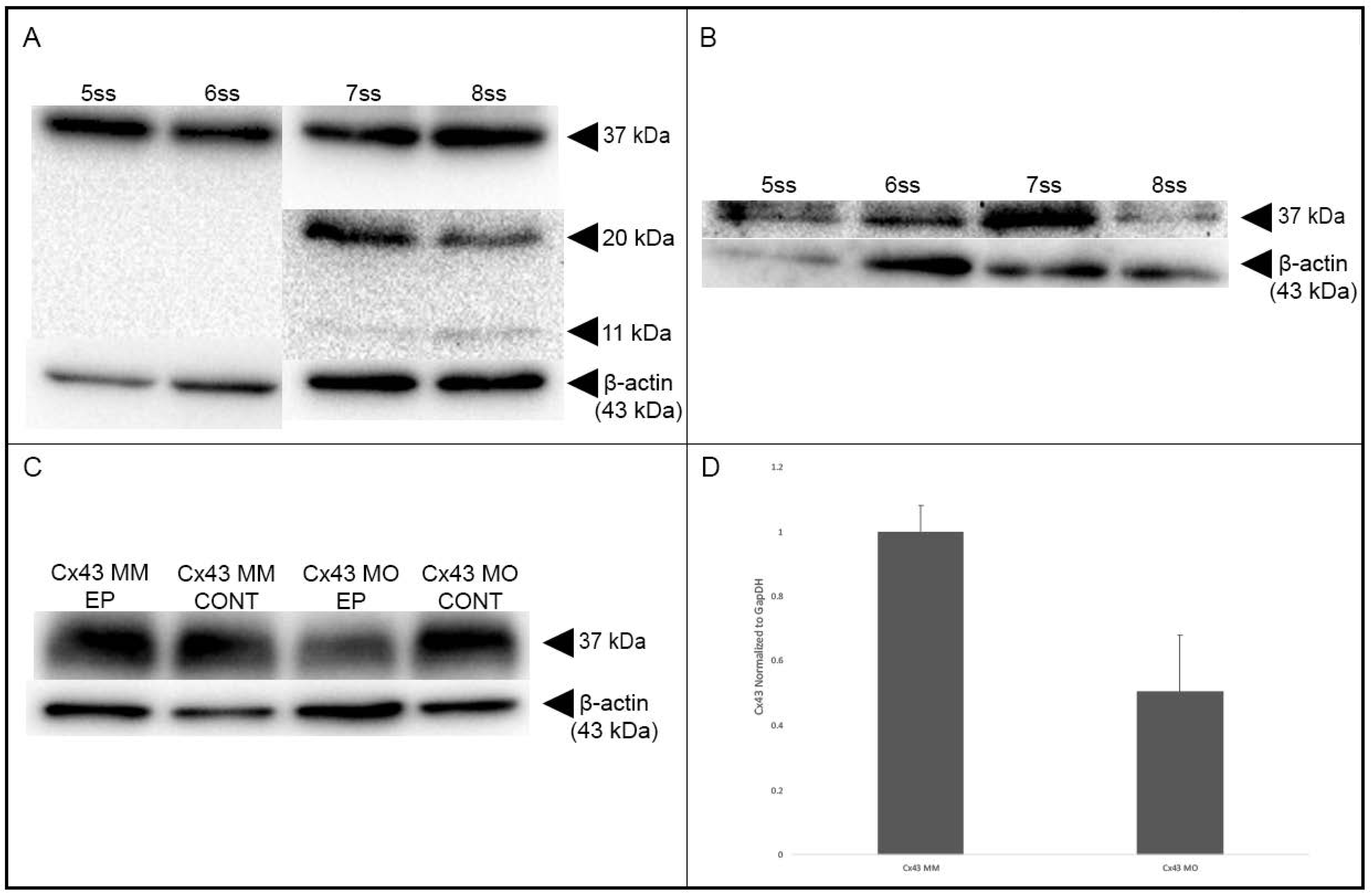
Immunoblotting for Connexin 43 validates the Connexin 43 antibodies and reveals different Connexin 43 isoforms during chick development. Immunoblotting results shown for antibodies directed against the C-terminus (A, C) or N-terminus (B) of Connexin 43 (Cx43). (A) The antibody directed against the C-terminus revealed the presence of full-length Cx43 protein (37 kDa) at all stages examined (5-8ss), while the 20 kDa and 11 kDa Cx43 isoforms are only present at the 7ss and 8ss. (B) The antibody directed against the N-terminus, which is distinct from the C-terminal epitope, does not detect the 20 kDa or 11 kDa isoforms of Cx43. Knockdown efficiency of the Cx43 MO was assessed as previously described (C. Y. Wu & Taneyhill, 2012; Shah *et al.*, 2017). Immunoblot analysis using the C-terminal antibody (C) reveals a 69% reduction (D) in Cx43 protein (Cx43 MO EP) as compared to the control MO-treated lysate (Cx43 MM EP) and associated contralateral controls (Cx43 MO CONT and Cx43 MM CONT).

### Premigratory and migratory cranial neural crest cells possess gap junctions that are dependent upon Cx43 for their function

To evaluate whether premigratory cranial neural crest cells form functional gap junctions, we examined the passive diffusion of Calcein dye, which only passes between cells via gap junctions (Davy *et al.*, 2006; Abbaci *et al.*, 2008), between premigratory neural crest cells *in vivo* containing the Cx43 control MO, which we validated by immunoblotting to have levels of Cx43 similar to the unelectroporated, contralateral control side of treated embryos (Fig. 1C, Cx43 MM EP vs. Cx43 MM CONT). In order to trace the spread of the Calcein between cells that had previously been electroporated with the Cx43 control MO, we co-injected a 1:1 solution of Calcein and Dextran, the latter being unable to pass through gap junctions due to its large size and is thus maintained in cells receiving the Calcein by injection (Ziambaras *et al.*, 1998). Results of these experiments demonstrated that, in under three hours, Calcein diffused to a small number of surrounding premigratory neural crest cells (Fig. 2A-A’’’, C-C’’’, E-E’’’, G-G’’’; arrowheads indicate dye spread; Supplemental Video 1). In embryos possessing premigratory cranial neural crest cells electroporated with the Cx43 MO, which leads to a 69% reduction in Cx43 protein levels by immunoblotting (Fig. 1C, Cx43 MO EP vs. Cx43 MO CONT; Fig. 1D), we noted no passage of Calcein to adjacent cells (Fig. 2B-B’’’, D-D’’’, F-F’’’, H-H’’’; Supplemental Video 2). Taken together, these results demonstrate that premigratory cranial neural crest cells form functional gap junctions and that Cx43 knockdown is sufficient to inhibit gap junction-based intercellular communication between them.

**Figure 2:**
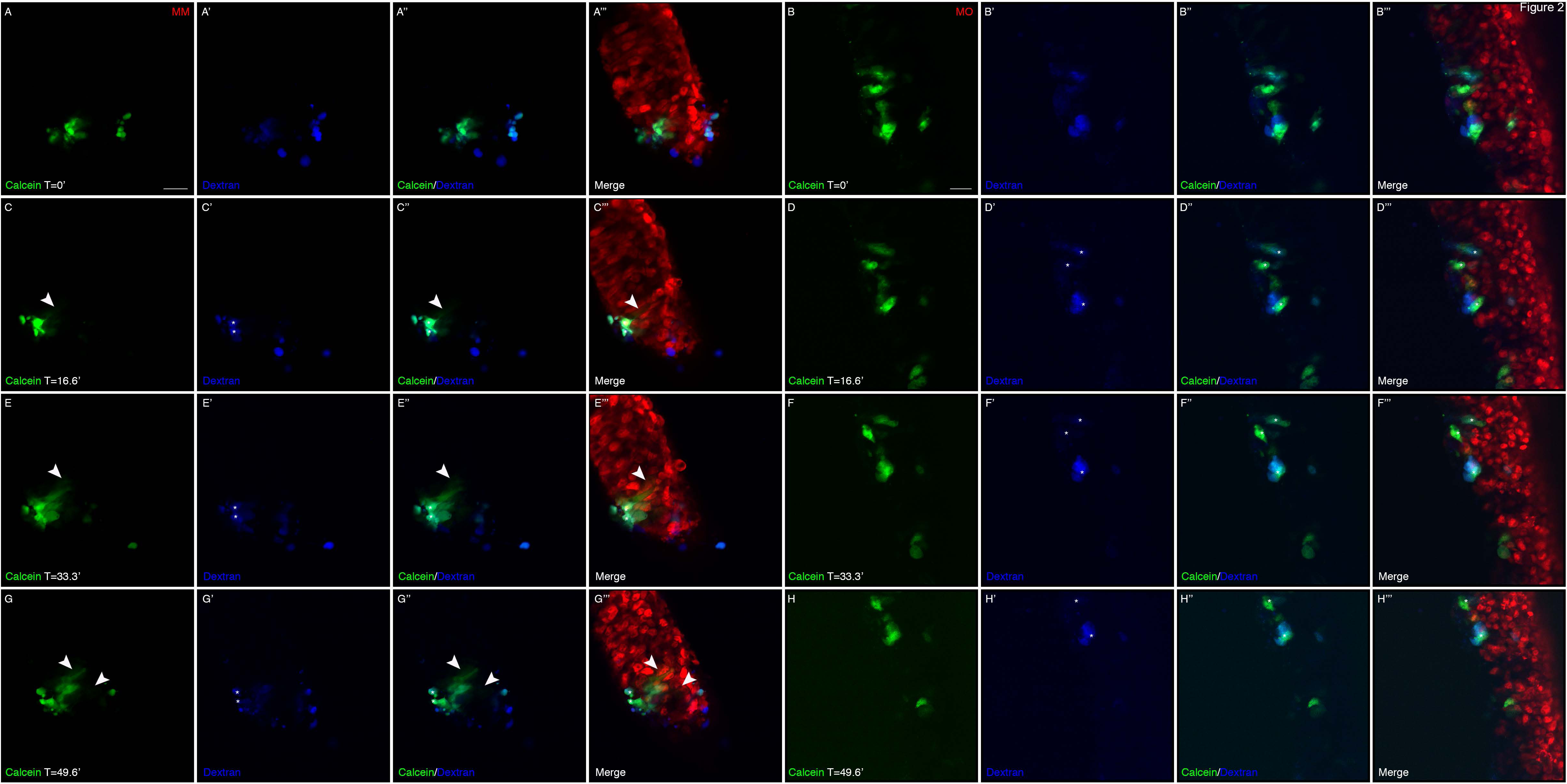
Morpholino-mediated knockdown of Cx43 abrogates gap junction function in chick premigratory cranial neural crest cells. Images of live neural folds showing diffusion of Calcein (green) from injected cells (identified with Dextran (blue)) to adjacent control MO (red)-electroporated premigratory neural crest cells (A-A’’’, C-C’’’, E-E’’’, G-G’’’; arrowheads) over a period of 50 minutes. No diffusion is observed with those cells possessing the Cx43 MO (red; B-B’’’, D-D’’’, F-F’’’, H-H’’’). Scale bars in (A) and (B) are applicable to their respective image sets and are 20 µm.

To determine whether migratory cranial neural crest cells also form functional gap junctions, we used an *ex vivo* culturing system to facilitate live imaging of dye passage, or gap junction function, between neural crest cells. To this end, dorsal neural folds (containing premigratory cranial neural crest cells) from either Cx43 control MO- or Cx43 MO-electroporated embryos were explanted into chamber slides and combined with dorsal neural folds previously treated with Calcein-AM for 30 minutes (Fig. 3A and K). After overnight incubation, during which time the neural folds attached to the dish and neural crest cells commenced EMT and migrated, we observed that control MO-treated migratory neural crest cells now possessed Calcein-AM within their cytoplasm due to their contact with adjacent Calcein-treated neural crest cells (Fig. 3C, arrows). We confirmed that these cells were, in fact, migratory neural crest cells by immunostaining fixed cultures with the HNK-1 antibody (Fig. 3B-E). We then repeated the experiment but performed live imaging at high magnification throughout the incubation period to observe dye transfer in real time. Through these assays, we showed that when a control MO-containing migratory neural crest cell extended a filopodia and made contact with an adjacent, Calcein-AM-positive migratory neural crest cell, the dye was taken into the control MO-containing neural crest cell in under four minutes (Fig. 3F-J, asterisk: Calcein contained within MO-positive cell; Supplemental Video 3).

**Figure 3:**
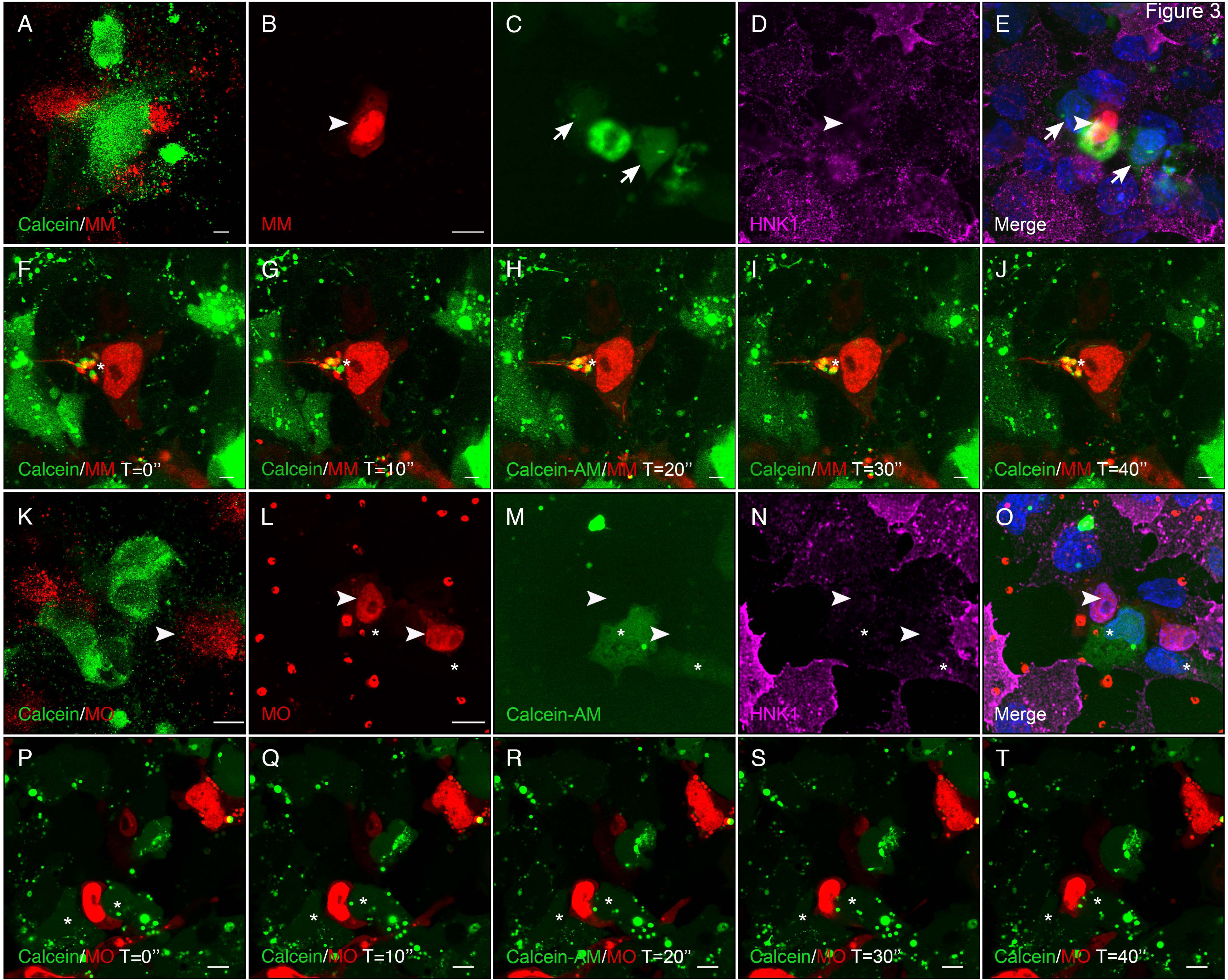
Morpholino-mediated knockdown of Cx43 abrogates gap junction function in chick migratory cranial neural crest cells. Images of live (A, F-K, P-T) or fixed and immunostained (B-E, L-O) neural crest cells showing diffusion of Calcein-AM dye (green) from Calcein-AM-treated wildtype (WT) neural crest cells to neural crest cells treated with either the Cx43 mismatch control (MM) or Cx43 morpholino (MO). When premigratory neural crest cells from MM-treated embryos were combined with Calcein-AM-treated WT premigratory neural crest cells (A-J), passage of Calcein-AM occurred between adjacent MM-positive neural crest cells, noted after fixation (B-E, arrowhead to arrow-labeled cells) and in live cultures (F-J, asterisk). No dye, however, moved to Cx43 MO-positive neural crest cells (K-O, arrowhead) while adjacent WT neural crest cells freely pass Calcein-AM (asterisk). This is more obvious during live imaging (P-T), where a MO-positive neural crest cell migrates between two adjacent cells and there is no transfer of Calcein-AM (asterisk). During live imaging, photos were acquired every 10 seconds (a total of 25 images were captured, but only the first five images are shown in stills; videos can be viewed in supplemental data). Scale bars in (A) and (K) are 100 µm; scale bars in (B-E) and (L-T) are 10 µm; and scale bars in (F-J) are 5 µm.

Next, we conducted the same fixed and live imaging assays described above but this time with Cx43 MO-containing neural crest cells. When Calcein-AM-positive neural folds were incubated with Cx43 MO-electroporated neural folds (Fig. 3K) and cultured to generate migratory neural crest cells from both populations, we did not detect a single cell in which dye transfer had occurred after immunostaining (Fig. 3L-O, arrowheads: MO-positive cells, asterisks: Calcein-positive cells). We confirmed these findings with live imaging, where we observed no dye transfer among Cx43 MO-containing neural crest cells migrating among adjacent, Calcein-AM-positive migratory neural crest cells over a period of five minutes of continuous imaging (Fig. 3P-T, asterisk: Calcein-positive cells; Supplemental Video 4). These results demonstrate that migratory neural crest cells also form gap junctions and that a reduction in Cx43 protein levels is sufficient to inhibit gap junction function in this cell population. Intriguingly, as premigratory neural crest cells were electroporated prior to EMT, these data also suggest that Cx43 knockdown is not sufficient to inhibit EMT in an *ex vivo* culture system.

### Cx43 knockdown reduces the size of the premigratory neural crest cell domain but does not inhibit neural crest cell EMT in vivo

To delineate the function of Cx43, and potentially gap junctions, on EMT independent of neural crest cell induction, we introduced the Cx43 control MO or Cx43 MO by unilateral electroporation at HH7^+^ to HH8^−^, three hours prior to the onset of EMT. Embryos were fixed and immunostained for a battery of premigratory and migratory neural crest cell markers (i.e., Cx43, Snail2, Sox10, Pax7, and E-cadherin) (Taneyhill *et al.*, 2007; Sauka-Spengler & Bronner-Fraser, 2008; Betancur *et al.*, 2010; Murdoch *et al.*, 2012; Lee *et al.*, 2013; Jourdeuil & Taneyhill, 2018). We also examined N-cadherin expression as it is a marker for the neural tube at this stage (Dady *et al.*, 2012; Lee *et al.*, 2013). When premigratory cranial neural crest cells were electroporated with the Cx43 control MO, we noted no change in any of the markers examined (Fig. 4A-A’’, C-C’’, E-E’’, G-G’’, I-I’’). As expected, electroporation of premigratory cranial neural crest cells with the Cx43 MO reduced the levels of Cx43 in both the dorsal neural folds and the neural tube (Fig. 4B-B’’, arrow). Surprisingly, however, reduction in the level of Cx43 resulted in a statistically significant decrease in the number of Pax7- (Fig. 4D-D’’, arrow; *p* = 0.007) and Snail2- (Fig. 4F-F’’, arrow; *p* = 0.0001) positive premigratory neural crest cells on the electroporated side compared to Cx43 control MO-electroporated premigratory neural crest cells (Table 1; all values were normalized to the contralateral control side prior to statistical analysis as per (Hutchins & Bronner, 2018)). Interestingly, the reduction in the number of Sox10-positive premigratory neural crest cells (Fig. 4H-H’’, arrow) was not statistically significant (*p* = 0.2). In addition, we saw no appreciable difference in levels and distribution of E-cadherin (not shown) and N-cadherin (compare Fig. 4I-I’’ to J-J’’). Importantly, this decreased domain size was not due to any changes in cell proliferation or cell death. Immunostaining with an antibody to phospho-histone H3 (Fig. 5A-B’’, asterisks in A’ and B’ indicate the premigratory neural crest cell population) or TUNEL assays (Fig. 5C-D’’, asterisks in C’ and D’ point to the premigratory neural crest cell population) showed no appreciable difference in cell proliferation or cell death, respectively, between control MO- and Cx43 MO-treated embryos. These results suggest that the reduction in the size of the premigratory neural crest cell domain may be due to an earlier effect on the cranial neural crest. Moreover, the absence of any change in the migratory neural crest population (Fig. 4D’, F’, H’, arrowheads) reveals that Cx43 knockdown is not sufficient to inhibit EMT *in vivo*, in keeping with our *ex vivo* explant culture data (Fig. 3).

**Figure 4:**
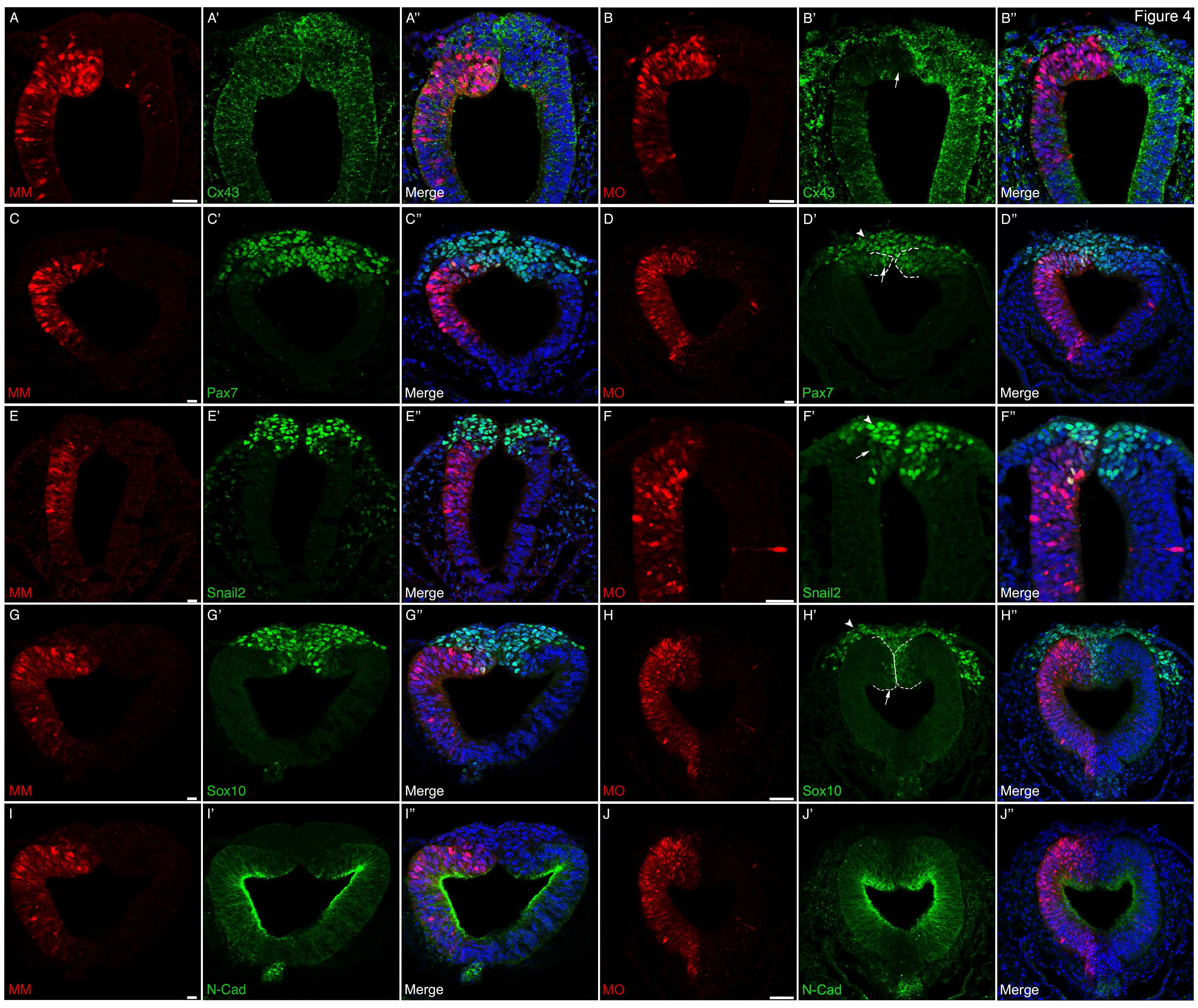
Morpholino-mediated knockdown of Cx43 decreases the number of premigratory neural crest cells in the absence of any changes in cell proliferation or cell death. Representative transverse sections taken through the midbrain and hindbrain of 5-9ss embryos after unilateral electroporation of the neural tube at the 3ss with either the mismatch control morpholino (MM) or Cx43 morpholino (MO) and immunostaining for various premigratory and/or migratory neural crest cell markers. In MM-treated embryos (A-A’’, C-C’’, E-E’’, G-G’’, I-I’’, K-K’’, M-M’’), there is no decrease in the level of Cx43 in the neural tube (A’) and no change in any premigratory and/or migratory neural crest cell markers (e.g., Pax7 (C’), Snail2 (E’), Sox10 (G’)). Similarly, there are no changes in N-cadherin (I-I’’), a marker of neural tube cells. In Cx43 MO-treated embryos (B-B’’, D-D’’, F-F’’, H-H’’, J-J’’, L-L’’, N-N’’), however, there is obvious knockdown of Cx43 in the neural tube (B’, arrow) and also a reduction in the premigratory neural crest cell population observed after Pax7 (D’, arrow), Snail2 (F’, arrow), and Sox10 (H’, arrow) immunostaining. Interestingly, other markers such as N-cadherin (J-J’’) and E-cadherin (not shown) are not affected. Cx43 knockdown does not seem to be required for EMT, as neural crest cells emerge from the neural tube (D’, F’, H’; arrowheads). Scale bars are 20 µm.

**Figure 5.**
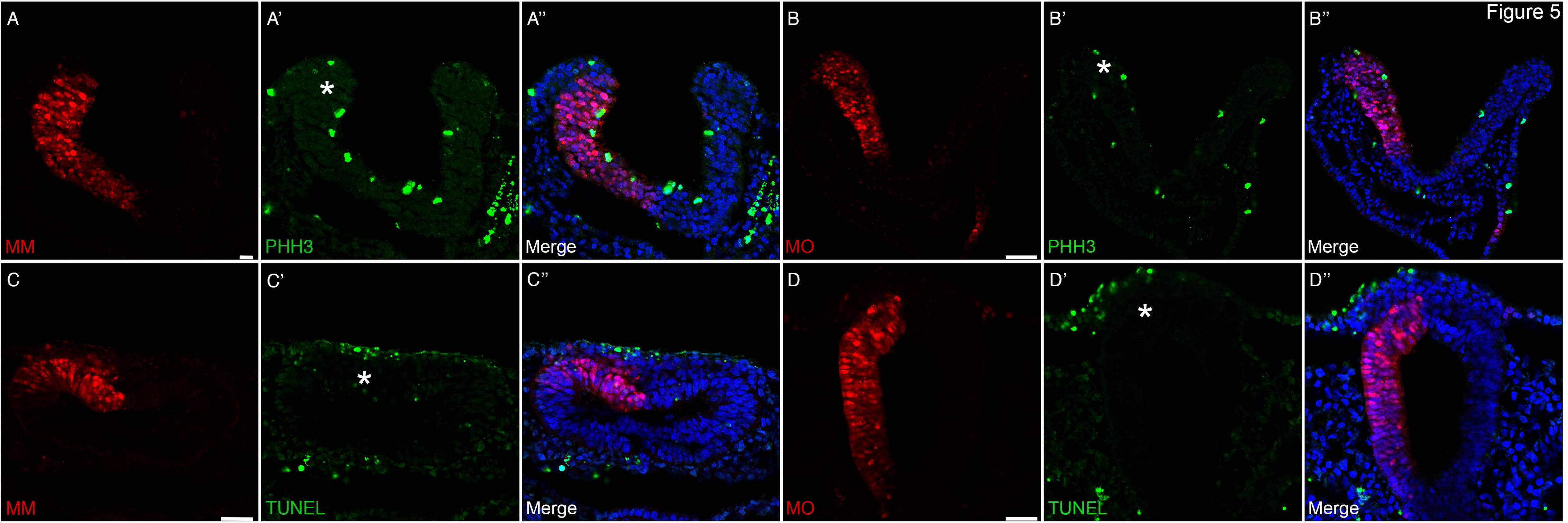
MO-mediated depletion of Cx43 does not affect cell proliferation or cell death in the neural tube and/or premigratory neural crest cell population. Representative transverse sections taken through the midbrain and hindbrain of 5-9ss embryos after unilateral electroporation of the neural tube at the 3ss with either the mismatch control morpholino (MM) or Cx43 morpholino (MO). Immunostaining for phospho-histone H3 (PHH3) revealed no difference in cell division between control MO-treated embryos (A-A’’) and Cx43 MO-treated embryos (B-B’’; asterisk indicates Cx43 MO-electroporated premigratory neural crest cell population in B’). Similarly, TUNEL staining revealed no difference in cell death between control MO-treated embryos (C-C’’) and Cx43 MO-treated embryos (D-D’’; asterisk indicates Cx43 MO-electroporated premigratory neural crest cell population in D’). Scale bars are 20 µm.

Given this change in the number of premigratory neural crest cells upon Cx43 knockdown, we next evaluated the function of Cx43-mediated gap junction communication in neural crest induction. To this end, we performed electroporations at HH4 to target the Cx43 control MO or the Cx43 MO into the epithelium comprising the non-neural ectoderm, neural plate border region (consisting of those cells that will give rise to neural crest and placode cells), and the neural ectoderm. While the Cx43 control MO-electroporated embryos developed normally, the Cx43 MO-electroporated embryos displayed significant malformations such that it was impossible to characterize any effects on neural crest induction (not shown). These data are not surprising as Cx43 is expressed from very early stages of mouse embryogenesis (5.5 days post coitum) through to the establishment of the neural crest (Yancey *et al.*, 1992; Lo *et al.*, 1997; Jourdeuil & Taneyhill, 2018; Kotini *et al.*, 2018) and is critical for a large number of developmental processes. Altogether, these results indicate an important role for Cx43 during normal chick morphogenetic movements and later in the establishment and/or maintenance of the premigratory cranial neural crest cell domain. Moreover, premigratory cranial neural crest cells do not require Cx43-dependent gap junction-mediated communication during EMT and early migration.

## DISCUSSION

### Premigratory and migratory neural crest cells form functional gap junctions

Previous research has focused on the expression of Cx43 and gap junction function in mouse cardiac neural crest cells, as *Cx43* knockout (KO) mice die from conotruncal heart malformations and defects associated with the outflow tract, a region of the heart that forms from cardiac neural crest cells (Ruangvoravat & Lo, 1992; Ewart *et al.*, 1997; Lo *et al.*, 1997; Huang *et al.*, 1998; Sullivan *et al.*, 1998; Xu *et al.*, 2001; Wei *et al.*, 2005; Xu *et al.*, 2006; Rhee *et al.*, 2009; Francis *et al.*, 2011). In mouse, *Cx43* is expressed in premigratory and migratory neural crest cells as well as those undergoing EMT (Ruangvoravat & Lo, 1992; Lo *et al.*, 1997). Furthermore, migratory neural crest cells form function gap junctions (Lo *et al.*, 1997). To further study Cx43 function, a transgenic mouse overexpressing *Cx43* (CMV43) (Ewart *et al.*, 1997) or a dominant-negative Cx43 fusion protein (Sullivan *et al.*, 1998) were generated. Cardiac neural crest cells in CMV43 mice possess an increased migration rate, with lower rates of migration observed for cardiac neural crest cells from the *Cx43* KO and dominant-negative Cx43 mouse (Huang *et al.*, 1998). In *Xenopus*, Cx43 MO-mediated knockdown only impacted neural crest migration, as none of the early premigratory neural crest markers were affected (Kotini *et al.*, 2018). In chick, we have shown that Cx43 is expressed in cranial neural crest cells from HH8^−^ through to HH17 during trigeminal ganglion assembly (Jourdeuil & Taneyhill, 2018). In all of these systems, however, gap junction presence and function in premigratory neural crest cells and those undergoing EMT remained poorly understood, until our studies herein.

Our results now show that gap junctions are present in both chick premigratory and migratory cranial neural crest cells and that their function is dependent on Cx43. An approximate 70% reduction in Cx43 levels within the premigratory and migratory cranial neural crest cell populations impeded dye transfer between adjacent neural crest cells. In the presence of Cx43, dye will readily pass from one cranial neural crest cell to an adjacent cell, but this is not observed upon Cx43 knockdown. The presence of Cx43-mediated gap junctions in the neural crest could imply a function for gap junction-based communication within the premigratory and migratory cranial neural crest cell populations, which we investigated in more detail in the chick embryo.

### Cx43 knockdown reduces the number of premigratory neural crest cells but does not affect neural crest cell EMT

After determining that gap junctions were formed between premigratory and migratory cranial neural crest cells, we next evaluated the function of Cx43 in the embryo during neural crest cell EMT. Strikingly, unilateral electroporation of the Cx43 MO at HH7^+^ and HH8^−^ (2-3ss) decreased the number of premigratory cranial neural crest cells but did not affect EMT. In addition, this reduction in the number of premigratory cranial neural crest cells was not due to increased apoptosis or decreased cell proliferation. This is different from what was observed in *Xenopus* embryos, where Cx43 MO-mediated reduction in the levels of Cx43 had no effect on the premigratory neural crest cell population (Kotini *et al.*, 2018). It should be noted that while a 70% reduction in Cx43 protein levels achieved by the Cx43 MO abrogated gap junction function (i.e., the ability of neural crest cells to transfer dye), it is possible that any residual Cx43 protein may be sufficient to mediate EMT. However, this would likely occur through a gap junction-independent function of Cx43 given the robust loss of intercellular dye passage. In addition, we cannot rule out that other connexins expressed in cranial neural crest cells may be capable of compensating for this reduction in Cx43, or function normally to mediate EMT by forming gap junctions. As the introduction of the Cx43 MO inhibited gap junction function and also decreased the number of premigratory cranial neural crest cells, it is possible that the normal function of Cx43-containing gap junctions is to permit the passive diffusion of an intercellular signal(s) required to maintain this cell population. A reduction in Cx43 could also affect hemichannel function, which allows small molecule and ion exchange to occur between the cytoplasm and the extracellular environment and has recently been shown to play a role in extracellular signaling and disease function (Nielsen *et al.*, 2012; Delmar *et al.*, 2018; Rovegno & Saez, 2018; Valdebenito *et al.*, 2018). It will be necessary in the future to develop techniques in the chick that will allow us to query hemichannel function in the cranial neural crest.

During the course of this study, we attempted to electroporate the presumptive neural crest cell population prior to neural crest cell induction (HH4) with either the Cx43 control or Cx43 MO to evaluate effects on this earlier event. While Cx43 control MO-electroporated embryos developed normally, the Cx43 MO had such a deleterious effect on morphogenesis that it was impossible to determine whether Cx43-mediated gap junctions are involved in neural crest induction (not shown). These data are not altogether surprising as Cx43 is expressed in a large number of tissues, including the ectoderm, neural crest, and neural tube; plays critical functions in intercellular communication, cytoskeletal organization, and signaling; and connexin dysfunction is associated with diseases that affect a variety of tissues and organs (Davy *et al.*, 2006; Dbouk *et al.*, 2009; Kanaporis *et al.*, 2011; Nielsen *et al.*, 2012; Laird, 2014; Delmar *et al.*, 2018; Jourdeuil & Taneyhill, 2018; Rovegno & Saez, 2018; Sorgen *et al.*, 2018; Valdebenito *et al.*, 2018; Cooreman *et al.*, 2019; Waning *et al.*, 2019; J. I. Wu & Wang, 2019). Therefore, techniques in the chick to knockdown or knockout Cx43 function specifically in the presumptive neural crest cell population at a well-defined time must be developed in order to evaluate the requirement for Cx43, and Cx43-mediated gap junction function, in the induction of cranial neural crest cells.

### Cx43 and gap junctions may play different and species-dependent roles in the neural crest

While we have identified the presence of Cx43-20k and Cx43-11k C-terminal isoforms in chick cranial neural crest cells during neural crest cell EMT and early stages of migration, their function at this time in the chick remains unknown. In *Xenopus*, MO-mediated knockdown of full-length Cx43 worked in a cell autonomous manner that specifically affected neural crest cell migration without interfering with neural crest cell induction. Furthermore, the Cx43-20k isoform directly regulates *N-cadherin* transcription in migratory neural crest cells through a BTF3/PolII complex (Kotini *et al.*, 2018). Interestingly, early studies to examine gap junction function in mouse migratory neural crest cells *ex vivo* by Lo and colleagues noted that carboxyfluorescein dye-filled cells, which had undergone gap junction coupling, expressed little to no N-cadherin at sites of cell-cell contact, contrary to what they expected to see in emerging neural crest cells (Lo *et al.*, 1997). Subsequent investigation into the role of N-cadherin during mouse cardiac neural crest cell migration revealed that N-cadherin is expressed at sites of cell-cell contact found between extended cell processes. and that N-cadherin is often co-localized, or closely apposed, to Cx43-positive gap junctions (Xu *et al.*, 2001). Interestingly, while N-cadherin deficiency also affected gap junction function in neural crest cells, Cx43 was not downregulated (Xu *et al.*, 2001), and, while N-cadherin appears to be required for Cx43 localization to the cell membrane (Wei *et al.*, 2005), Cx43 and N-cadherin appear to have separable roles in cardiac neural crest cell migration (Xu *et al.*, 2001). Additionally, these studies do not suggest a role for Cx43 in *N-cadherin* transcription, highlighting the diverse function of Cx43 in different species. As Cx43 interacts differently with N-cadherin in both *Xenopus* and mouse neural crest cells, it is therefore likely that Cx43/N-cadherin interactions may differ among species.

In the chick, N-cadherin is expressed in the neural tube but is absent in premigratory neural crest cells prior to EMT (Hatta & Takeichi, 1986; Nakagawa & Takeichi, 1998; Dady *et al.*, 2012; Jourdeuil & Taneyhill, 2018). Moreover, if N-cadherin is ectopically expressed in trunk premigratory neural crest cells, no neural crest cells undergo EMT (Nakagawa & Takeichi, 1998). Migratory neural crest cells in the chick, however, express Cadherin-7 (Nakagawa & Takeichi, 1998). Therefore, the function of the Cx43-20k isoform during early neural crest cell migration in the chick remains to be elucidated as does whether Cadherin-7 can interact with Cx43 in a manner similar to what has been described for N-cadherin in mouse.

Remarkably, our study shows a significant reduction in the size of the premigratory cranial neural crest cell population after Cx43 knockdown, which has not been described in other species after loss of Cx43. The reduction in the size of the premigratory cranial neural crest may be due to the loss of gap junction function or loss of a required gap junction-independent function of Cx43. To unravel this, it will be necessary to target earlier stages during the induction and specification of the neural crest; however, our attempts were unsuccessful due to the apparent requirement for Cx43 in proper morphogenetic movements during early chick development.

These data also suggest, however, that connexins (and potentially gap junctions), may play different roles in neural crest cell induction, specification, EMT, and migration in different species or different subpopulations of the neural crest (i.e., cardiac versus cranial). As there is robust expression of full-length Cx43 at these stages in chick, and reduction in Cx43 inhibits gap junction function, gap junctions may play a role in coordinating cell-cell communication during the establishment of the neural crest cell population. The function of Cx43-mediated gap junctions during neural crest cell migration in chick is currently unknown and is an area of current study. During EMT, however, it is likely that the function of Cx43 is gap junction-independent, possibly involving Cx43-mediated cytoskeleton rearrangements as identified in mouse cardiac neural crest cells, which will be followed up in a future study.

## CONCLUSIONS

These data reveal that both premigratory and migratory cranial neural crest cells possess functional gap junctions in chick and that the function of these gap junctions depends, in part, on Cx43. Furthermore, reduction of Cx43 protein levels between HH7^+^ and HH8^−^ decreases the number of Pax7- and Snail2-positive premigratory neural crest cells but is insufficient to block EMT. This suggests that gap junctions may be needed to maintain the size of the premigratory neural crest cell population but are not required for EMT. As we are unable to achieve complete knockdown of Cx43, however, we cannot determine whether residual Cx43 is serving a non-gap junction-related function and therefore allowing EMT to progress normally. Further study will be required to determine such a role for Cx43 during cranial neural crest cell EMT, as initial experiments to decipher the function of Cx43 during early neural crest cell induction were not fruitful due to large-scale developmental abnormalities occurring upon Cx43 knockdown at this stage. Interestingly, we also identified multiple C-terminal isoforms of Cx43 (Cx43-20k and Cx43-11k) in chick cranial neural crest cells during EMT and early migration. A recent paper in *Xenopus* demonstrated that the Cx43-20k isoform is a transcriptional regulator of *N-cadherin*, a gene required for neural crest migration in *Xenopus*. Chick cranial neural crest cells do not require N-cadherin to mediate their early migration, however, suggesting that connexins may play distinct roles in different species. Taken together, these data suggest that connexins, and gap junctions, may be critical for intercellular communication to maintain the appropriate size of the premigratory cranial neural crest cell domain prior to these cells undergoing EMT and migrating to properly pattern the vertebrate embryo.

## ACKNOWLEDGMENTS

This research was supported by grants from the National Institutes of Health (NIH R01DE024217, L.A.T.) and the American Cancer Society (RSG-15-023-01-CSM, L.A.T.). The authors would also like to thank Ms. Irina Kolesnik, Ms. Sophia Liu, and Ms. Caroline Halmi for excellent technical assistance.

