## Supplemental Video Legends for "Connexin 43 impacts the chick premigratory cranial neural crest cell population without affecting the neural crest cell epithelial-to-mesenchymal transition"

**SUPPLEMENTAL VIDEO FILES**

**Supplemental Video 1: Cx43 control MO electroporation does not affect gap junction function in the premigratory cranial neural crest cell population *in vivo***. Images were acquired every approximately 16 minutes for a total of 50 minutes.

**Supplemental Video 2: Cx43 MO-mediated knockdown of Cx43 abrogates gap junction function in the premigratory cranial neural crest cell population *in vivo***. Images were acquired every approximately 16 minutes for a total of 50 minutes.

**Supplemental Video 3: Cx43 control MO electroporation does not affect gap junction function in the migratory cranial neural crest cell population in an *ex vivo* culture assay**. Images were acquired every 10 seconds for a total of approximately 250 seconds.

**Supplemental Video 4: Cx43 MO-mediated knockdown of Cx43 abrogates gap junction function in migratory neural crest cells in an *ex vivo* culture assay**. Images were acquired every 10 seconds for a total of approximately 250 seconds.
